# Presumptive detection of *Giardia lamblia* cysts in borehole water in Gombe metropolis, north-eastern Nigeria: a two-round cross-sectional study with implications for water safety

**DOI:** 10.64898/2026.10.25.761622

**Authors:** Anas Ayuba, Esther Kela

**Affiliations:** Department of Science Laboratory Technology, Gombe State University, Tudun Wada, Gombe State, Nigeria

**Author notes:** Corresponding author: Esther Kela, Anas Ayuba.

**Keywords:** *Giardia lamblia*, borehole water, groundwater, waterborne parasites, water safety, Nigeria

## Abstract

*Giardia lamblia* (syn. *G. duodenalis, G. intestinalis*) is a flagellated intestinal protozoan and a common cause of waterborne diarrhoeal disease. Its environmentally resistant cysts can persist in untreated groundwater, yet boreholes in Nigeria are widely assumed to be safe and are rarely screened for protozoan parasites. We examined borehole water from ten quarters of Gombe metropolis (Malam Inna, Tudun Wada, Pantami, Orji, Tumfure, Bolari, Jekadafari, Nassarawo, Kumbiya Kumbiya and Kagarawal) in a cross-sectional study with two sampling rounds two weeks apart. One 15 L sample was collected per site per round (20 samples in total), concentrated by sedimentation and centrifugation, and examined as unstained normal-saline wet mounts by light microscopy. Samples were scored positive when at least one cyst with morphology consistent with *Giardia* was seen. Two of 10 samples (20%) were positive in each round; overall, 4 of 20 samples (20%; 95% Wilson CI 8–42%) were positive. Three of ten sites were positive at least once: Jekadafari in both rounds, indicating repeated detection, and Tumfure and Pantami in one round each. Positivity did not differ between rounds (Fisher’s exact test, *p* = 1.000). Because identification relied on morphology alone, without staining, immunofluorescence, PCR or viability testing, and because pH, turbidity and temperature were not measured on site, these results are presumptive detections from a small, non-random sample of boreholes. They nevertheless indicate that the underground origin of borehole water does not by itself guarantee freedom from parasitic contamination. We recommend routine parasitological monitoring, wellhead protection and household water treatment, and molecular confirmation (PCR and genotyping) in follow-up work.

## 1 Introduction

Access to safe drinking water remains a major public health challenge in sub-Saharan Africa, where waterborne disease contributes substantially to illness and death, particularly among children (WHO 2022). In many urban and peri-urban settings, boreholes are a principal alternative to inconsistent piped supply and are widely perceived as safe because groundwater is assumed to be naturally filtered. In practice, borehole water is typically consumed without any disinfection barrier.

*Giardia lamblia* infects the small intestine of humans and many other mammals and causes giardiasis, characterised by diarrhoea, abdominal cramps, bloating, nausea and malabsorption (Adam 2001). The motile trophozoite colonises the intestine, whereas the cyst (about 8–12 µm long, with a thick refractile wall and four nuclei when mature) is excreted in faeces and transmits infection through contaminated water and food (Adam 2001). The infectious dose is low, with infection established by as few as 10–100 cysts (Adam 2001; Feng and Xiao 2011). Cysts survive for weeks to months in cool water (deRegnier et al. 1989; Olson et al. 1999) and are more resistant to chlorine than enteric bacteria (Rice et al. 1982; LeChevallier et al. 1991), although they are susceptible to ultraviolet irradiation, boiling and appropriately designed filtration (Hijnen et al. 2006; WHO 2017). Waterborne outbreaks of giardiasis are well documented, including in settings where monitoring did not specifically look for protozoa (Karanis et al. 2007; Nygård et al. 2006). Giardiasis was added to the WHO Neglected Diseases Initiative in 2004 (Savioli et al. 2006).

Groundwater protection is not absolute. It depends on water-table depth, overlying geology, preferential flow paths and the integrity of borehole casing and wellhead (WHO 2017). On-site sanitation is a major source of faecal loading to groundwater in many low- and middle-income settings, with risk depending on latrine density, distance from the water source and soil and aquifer characteristics (Graham and Polizzotto 2013). Where cysts reach an untreated borehole, they can be delivered directly to consumers (Betancourt and Rose 2004). Nigerian studies of parasitic contamination of drinking-water sources generally find boreholes less contaminated than surface water or hand-dug wells, but not consistently free of parasites. *Giardia lamblia* has been recovered from boreholes in Kogi and Ondo States, whereas no parasites were found in boreholes in Dutsin-Ma, Katsina State (Iyaji et al. 2016; Simon-Oke et al. 2020; Abdullahi et al. 2018). A meta-analysis reported a pooled *Giardia* prevalence of 15.4% in Nigerian waterbody samples examined by microscopy (Tawana et al. 2023). Most Nigerian studies report parasites as a group, examine small volumes (for example 100 mL; Simon-Oke et al. 2020), and sample each site once. Routine monitoring focuses on bacteriological indicators, which do not reliably predict the presence of protozoan cysts (WHO 2017).

To our knowledge, there are few or no published data on *Giardia* in borehole water in Gombe metropolis, where many residents rely on untreated boreholes. We therefore determined the occurrence of *Giardia* cysts in borehole water from ten quarters of Gombe metropolis using large-volume (15 L) samples collected on two occasions two weeks apart, to describe overall positivity and to distinguish repeated from transient detection at the same sites.

## 2 Materials and Methods

### Study area and design

The study was conducted in ten quarters of Gombe metropolis, Gombe State, Nigeria: Malam Inna, Tudun Wada, Pantami, Orji, Tumfure, Bolari, Jekadafari, Nassarawo, Kumbiya Kumbiya and Kagarawal (Fig. 1). One designated borehole serving as a public or domestic water source was sampled in each quarter. This was a cross-sectional, comparative field study with two sampling rounds two weeks apart at the same ten sites. The sites were designated rather than randomly selected, so the results describe the sampled boreholes and not all boreholes in the metropolis.

**Figure 1:**
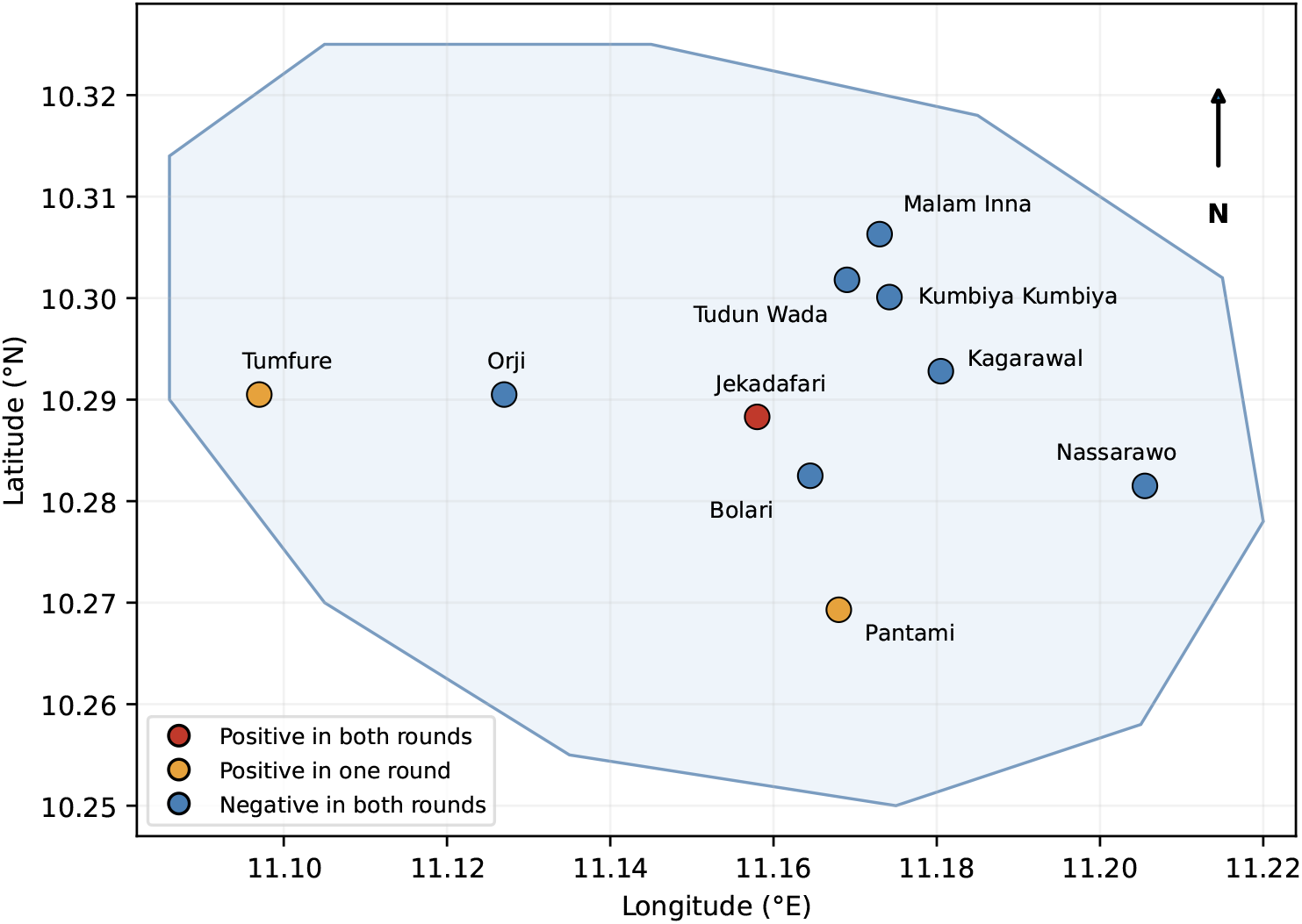
Schematic map of the ten sampling quarters in Gombe metropolis, coloured by the number of positive rounds. Markers show approximate quarter locations and not the exact positions of the sampled boreholes; the outline is schematic and not an administrative boundary.

### Sample collection

A 15 L water sample was collected from each borehole in each round, giving 20 samples (2 rounds *×* 10 sites). Samples were collected in clean sterile containers, sealed, labelled with site, date and round, and transported to the laboratory for processing. Collectors wore gloves and personal protective equipment. Sampling was carried out in August 2026 with the permission of the borehole owners, and the two rounds were two weeks apart.

### Laboratory examination

Each 15 L sample was left to stand for 48 h so that suspended material and heavier organisms could sediment. The sediment was recovered and centrifuged at 3000 rpm for 10 min to concentrate any cysts. One slide was prepared for each sample (20 slides in total): one drop of the concentrated sediment was mixed on a clean glass slide with one drop of normal saline, covered with a cover slip, and examined under a compound light microscope by a trained observer. No stain was applied. A sample was scored positive if at least one oval to ellipsoidal, refractile body with a distinct cyst wall, morphologically consistent with a *Giardia* cyst, was observed, and negative if none was seen. This sedimentation–centrifugation– saline wet-mount approach follows the general principle used in comparable Nigerian water-parasitology studies (Oyibo et al. 2018).

### Statistical analysis

Positivity was summarised as counts and percentages, with 95% Wilson confidence intervals. Because each round had *n* = 10 and some expected cell counts were below 5, positivity in rounds 1 and 2 was compared with Fisher’s exact test and with the chi-square test with Yates’ continuity correction (significance at *p <* 0.05). Because the same ten sites were sampled in both rounds, an exact McNemar test on the paired site-level results was also performed.

## 3 Results

### Overall positivity

Twenty 15 L samples were examined. Two of 10 samples (20%) were positive in round 1 and two of 10 (20%) in round 2, giving 4 of 20 positive samples overall (20%; 95% CI 8–42%) and 16 negative (80%) (Table 1; Fig. 2A). Because the same sites were sampled twice, the 20 samples are not independent observations.

**Figure 2:**
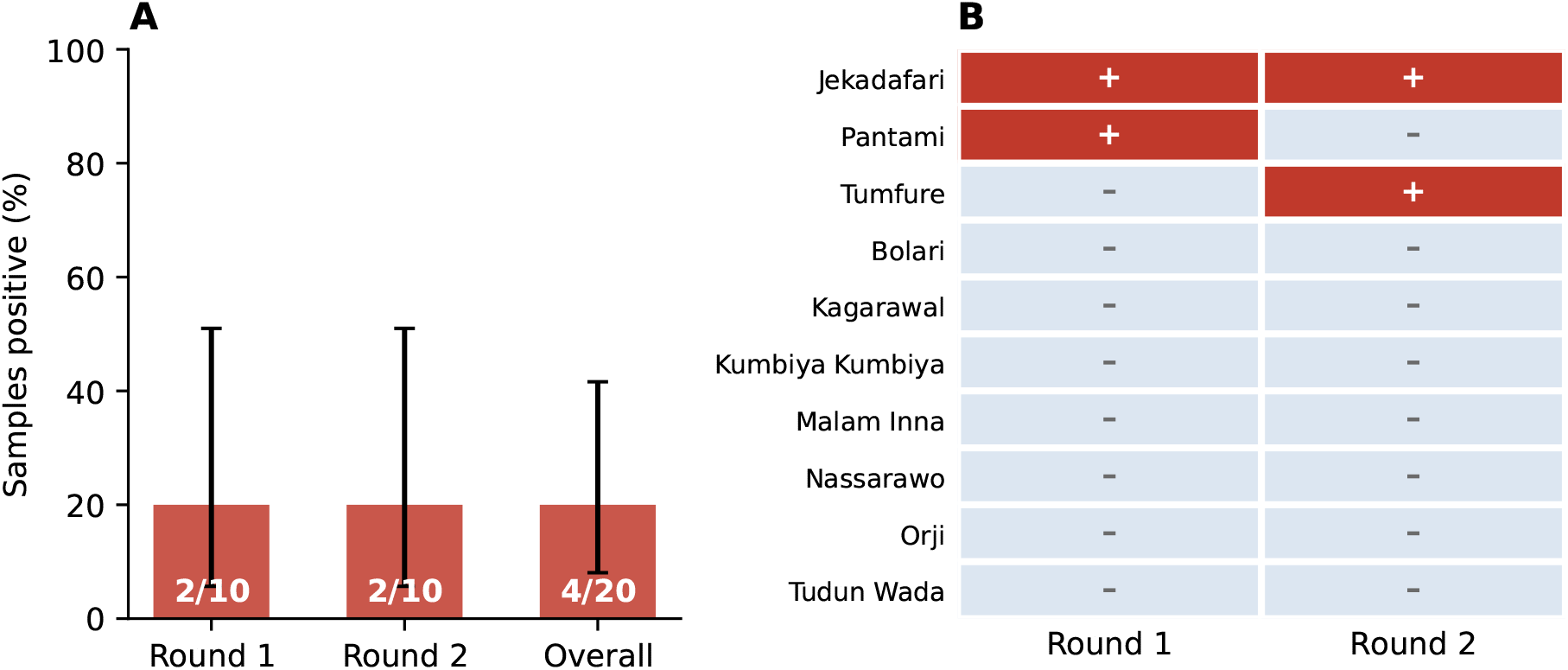
Microscopy results. (A) Percentage of samples positive in each round and overall, with 95% Wilson confidence intervals; numbers show positive/total samples. (B) Result for each site and round (+, positive; *−*, negative), ordered by number of positive rounds.

**Table 1:** Detection of *Giardia* cysts by wet-mount microscopy, by sampling round.

| Round | Samples | Positive | Negative | % positive | 95% CI |
| --- | --- | --- | --- | --- | --- |
| 1 | 10 | 2 | 8 | 20 | 6–51 |
| 2 | 10 | 2 | 8 | 20 | 6–51 |
| Total | 20 | 4 | 16 | 20 | 8–42 |

### Site-level results

Three of the ten sites (30%; 95% CI 11–60%) were positive at least once (Table 2; Fig. 2B). Jekadafari was positive in both rounds, indicating repeated detection at that borehole across the two-week interval. Pantami was positive in round 1 only and Tumfure in round 2 only, and the other seven sites (Malam Inna, Tudun Wada, Orji, Bolari, Nassarawo, Kumbiya Kumbiya and Kagarawal) were negative in both rounds.

**Table 2:** Detection of *Giardia* cysts by site and sampling round. +, cysts morphologically consistent with *Giardia* observed; *−*, none observed.

| Quarter | Round 1 | Round 2 |
| --- | --- | --- |
| Jekadafari | + | + |
| Pantami | + | – |
| Tumfure | – | + |
| Malam Inna | – | – |
| Tudun Wada | – | – |
| Orji | – | – |
| Bolari | – | – |
| Nassarawo | – | – |
| Kumbiya Kumbiya | – | – |
| Kagarawal | – | – |

### Comparison between rounds

Positivity did not differ between rounds (two of 10 in each; Fisher’s exact test *p* = 1.000; chi-square with Yates’ correction *χ*^2^ = 0.00, *p* = 1.000). One site was positive only in round 1 (Pantami) and one only in round 2 (Tumfure), and the exact McNemar test on these discordant pairs gave *p* = 1.000. Overall positivity was therefore identical in the two rounds, although apart from Jekadafari the positive sites differed between them.

## 4 Discussion

Cysts consistent with *Giardia lamblia* were detected in 4 of 20 (20%) borehole water samples from Gombe metropolis, at 3 of 10 sampled boreholes at least once, and in both rounds at one site (Jekadafari). All detections are presumptive, because they rest on cyst morphology in unstained saline wet mounts. With that caveat, the results indicate that the underground origin of borehole water does not guarantee freedom from parasitic contamination in this setting. This agrees with earlier reports that coliform testing alone can miss protozoan cysts and that unprotected boreholes can deliver them directly to consumers (Betancourt and Rose 2004; Karanis et al. 2007).

The lack of a difference between rounds (all *p* = 1.000) should be read cautiously: with ten samples per round the study had little power to detect a change. The site-level pattern is more informative. Repeated detection at Jekadafari is compatible with a persistent source of contamination, such as borehole construction, siting near pit latrines or other sources of faecal material, or seasonal changes in groundwater flow (Graham and Polizzotto 2013), whereas single-round detections at Pantami (round 1) and Tumfure (round 2) may reflect intermittent contamination or limited sensitivity of microscopy at low cyst concentrations. Neither explanation can be tested with our data. With only two rounds two weeks apart, we cannot distinguish a persistent source from chance repeated detection, nor describe seasonal variation, which has been associated with parasite occurrence in Nigerian water sources (Simon-Oke et al. 2020).

Overall positivity (20%; 95% CI 8–42%) is close to the 15.4% pooled prevalence for Nigerian waterbody samples (Tawana et al. 2023) and the 13.3% reported for borehole samples in Okura District, Kogi State (Iyaji et al. 2016), and the confidence interval is wide enough to include both. Boreholes were the least contaminated source in Akure and had no detected parasites in Dutsin-Ma (Simon-Oke et al. 2020; Abdullahi et al. 2018). Differences between studies may reflect local aquifer protection, borehole construction and proximity to on-site sanitation. We examined 15 L per sample, whereas some earlier studies examined far smaller volumes (100 mL in Akure; Simon-Oke et al. 2020), which increases the chance of recovering the few cysts that may be present. Conversely, identification by morphology alone means that false-positive identifications, for example of yeasts, algae, pollen or other protozoan cysts, cannot be excluded (Garcia 2016; Xiao and Fayer 2008). Comparisons across studies are also limited by differences in sample volume, methods and the sources sampled, and our design (one designated borehole per quarter, ten per round) does not support estimating the prevalence of contamination across the metropolis.

This study has important limitations. (i) Identification relied on morphology in unstained saline wet mounts; nuclei are less distinct without stain, microscopy is less sensitive than antigen-based or molecular methods (Sikiru et al. 2020), and microscopy cannot show viability, infectivity, species or assemblage (Garcia 2016; Xiao and Fayer 2008). Assemblages A and B infect humans, whereas C to H are largely animal-adapted (Cacciò and Ryan 2008; Feng and Xiao 2011), so the public health significance of the cysts detected is not yet established. (ii) pH, turbidity and temperature could not be measured on site because of lack of equipment, and WHO guidance recommends on-site measurement because these parameters change during storage and transport (WHO 2017); we therefore cannot relate detection to water quality or explain why cysts were found at particular sites. (iii) Sampling was limited to ten designated boreholes on two occasions, and faecal indicator bacteria, borehole depth and construction, and distance to latrines were not recorded. (iv) No positive or negative controls or recovery efficiency are reported, and the volume of sediment examined per sample was small compared with the volume sampled. Standard reference methods, such as US EPA Method 1623.1 and ISO 15553, use large-volume filtration, immunomagnetic separation and immunofluorescence (US EPA 2012; ISO 2006), and were not available to us.

Despite these limitations, the findings support treating untreated borehole water in Gombe metropolis as potentially at risk of protozoan contamination until shown otherwise. Because cysts are relatively resistant to chlorine, household measures such as boiling or appropriately designed filtration are more dependable than chlorination alone (WHO 2017). We recommend: (1) including protozoan parasites, not only bacteriological indicators, in routine borehole monitoring; (2) giving Jekadafari priority for investigation, with review of wellhead protection, casing integrity and distance from latrines and other sources of contamination; (3) equipping field teams with portable meters for on-site pH, turbidity and temperature; (4) promoting household water treatment; and (5) confirming and characterising detections with staining, immunofluorescence or PCR and genotyping, across more sites and sampling rounds, ideally covering both wet and dry seasons.

## 5 Conclusions

Cysts morphologically consistent with *Giardia lamblia* were found in 4 of 20 borehole water samples (20%) from ten quarters of Gombe metropolis, at 3 of 10 sites at least once, and in both rounds at Jekadafari. Although these presumptive microscopy-based findings need molecular confirmation, they show that borehole water in this setting should not be assumed to be free of parasites and that routine parasitological monitoring and household treatment are warranted.

## Declarations

### Author contributions

A.A. conceived and designed the study, collected samples, performed the laboratory work and analysis, and wrote the manuscript. E.K. supervised the study, advised on design and interpretation, and reviewed and edited the manuscript. Both authors approved the final version.

### Ethics statement

The study analysed environmental water samples and involved no human participants, animals or personal data. Water was sampled with the permission of the borehole owners.

### Data availability

All data generated in this study are included in the article (Tables 1 and 2).

## Acknowledgements

The authors thank Mr. Sani Jibir Waziri and Assoc. Prof. Buhari Magaji for their support, and the lecturers, laboratory technologists, colleagues and friends in the Department of Science Laboratory Technology, Gombe State University, who contributed to this work. This preprint is based on the first author’s B.Sc. final-year project (UG20/SCSL/1061), Gombe State University, 2026.

